# Tailoring 3D cell culture templates with common hydrogels

**DOI:** 10.1101/370882

**Authors:** Aurélien Pasturel, Pierre-Olivier Strale, Vincent Studer

**Author notes:** Correspondence and requests for materials should be addressed to V.S.

## Abstract

Hydrogels are the simplest and most widespread 3D cell culture materials, but turning these homogeneous substrates into biomimetic templates proves difficult. To this end, we introduce a generic solution compatible with the most biologically relevant and often frail materials. Here we take control of the chemical environment driving radical reactions to craft common gels with patterned light. In a simple microreactor, we harness the well-known inhibition of radicals by oxygen to enable topographical photopolymerization. Strikingly, by sustaining an oxygen rich environment, we can also induce hydrogel photo-scission, which turns out to be a powerful and generic subtractive manufacturing method. We finally introduce a flexible patterned functionalization protocol based on available photo-linkers. Using these tools on the most popular hydrogels, we tailored soft templates where cells grow or self-organize into standardized structures. The platform we describe has the potential to set a standard in future 3D cell culture experiments.

**Significance:** Light based, engineering of hydrogel should not be solely restricted to chemically modified materials. Indeed, many researchers rely on common hydrogels to organ-ize cells but lack the structuration and functionalization methods to develop standardized in vitro models with human-like properties. We unlock this limitation by providing a scalable toolbox that operates on widespread generic hydrogels or hydrogel blends such as PEG, Agar, or Matrigel. We crafted tailored 3D cell culture templates with these materials, growing and self-organizing cell lines and primary neurons in a controlled manner. More largely, we encourage all engineers with a knack for hydrogels but who are not experts in organic chemistry, to adopt our toolbox.

---

Hydrogels are commonplace in 3D cell culture i.e. the art of organ-izing cells by cultivating them in configurations that more closely mimic the in-vivo environment(1). Whether to fill scaffolds with cells(2), exploit cell self-organization(3–5), or both(6), their high hydric content, modulable mechanical properties(3) and access to chemical functionalization(7–9) make hydrogels a staple for most applications.

With the help of light, the ability to shape(10), cleave(11, 12) or functionalize(8, 9) these materials has improved in tandem with the idea that standardized organoid models will emerge from spatially defined micro-niches(6, 13). To this end, an array of light-based devices and photo-controllable hydrogels have been designed with the intent of turning these homogeneous matrixes into heterogeneous micro-environments(8, 11). Alternatively, laser-scanning photoablation has been used to degrade natural matrixes(12) like Matrigel or agar-agar which are commonplace(5, 14) but lack photo-responsive moieties.

Unfortunately, despite their respective successes, these technologies cannot favor widespread adoption as they either lack the required flexibility, accessibility or speed. To keep biology at the center of focus, an ideal toolbox should alleviate labor-intensive developments in chemistry, hardware and 3D modelling thus simplifying the conception of microniches out of common place materials.

In this report, we introduce a setup consisting of a commercially available widefield patterned light illuminator and oxygen permeable PDMS micro-reactors. This platform can spatially control the kinetics of light-based reactions by locally tuning the photon flux over the entire insolated field.

This flexible solution performs: (i) additive manufacturing of photopolymerizable hydrogels with control over thickness (ii) subtractive manufacturing of many gels such as PEG, Agar, Matrigel or poly-acrylamide (iii) decoration (patterned functionalization) of hydrogels.

Queuing and combining the above capabilities, we turned common hydrogels into topographically and chemically heterogeneous micro-environments. Cells were then seeded onto those templates, growing or self-organizing into various designs: standardized spheroids, patterned cells on hydrogels and colonized perfusable channels.

In short, a flexible platform, common materials and a simple workflow open an avenue of hydrogel engineering for 3D cell culture.

## Results

### A set-up to control the photo-initiator’s chemical context

To engineer hydrogels with light, our platform makes extensive use of a benzophenone-based photo-initiator. Upon irradiation, benzophenone forms highly reactive radicals that can induce polymer crosslinking, degradation and functionalization depending on its immediate chemical environment (gas, pre-polymers, photo-linker)(15). To harness this sensitivity towards the environment, we developed modular microreactors into which reagents are injected and gas-gradients can be generated, while photochemical reactions occur (Fig.1A).

**Fig. 1.**
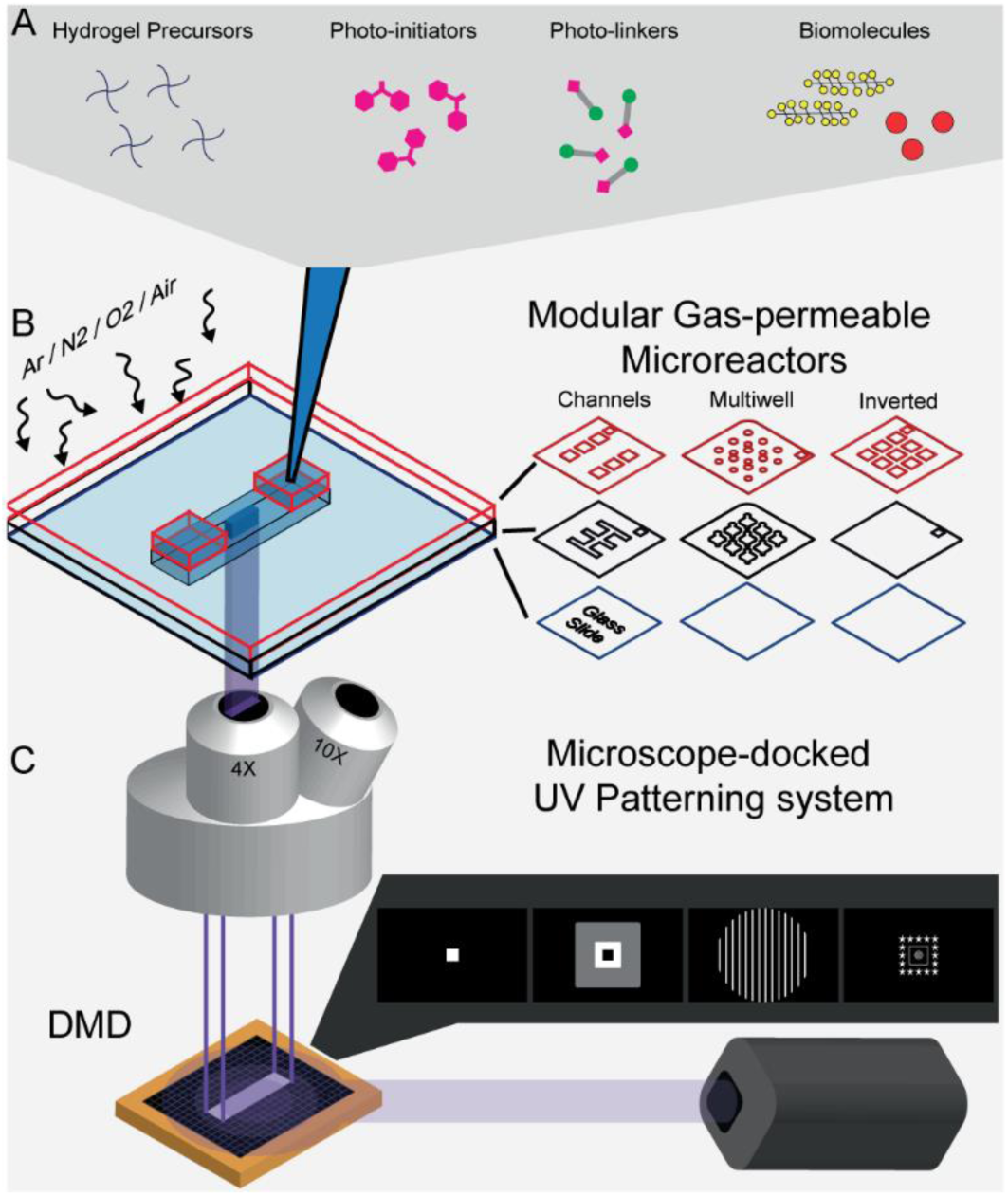
A flexible set-up to control reagents, gas and photon distribution. (A) Schematics of the potential reagents used in the platform with hydrogel precursors (blue 4-branched stars), photo-initiator (pink), photo-linkers (pink and green), biomolecules (yellow, red). (B) Schematics of the gas permeable PDMS microreactors used in this paper showing the upper (red) and lower (black) PDMS stencil designs stacked onto a glass slide (blue). (C) Schematics of the DMD-based (gold) UV projection set-up with microscope objectives (gray), projected patterns and 375nm expanded and collimated laser beam (purple).

These microreactors are made of two gas-permeable PDMS stencils(16) stacked onto a glass microscopy slide enabling oxygen gradients as well as changes in atmospheric content. These foils of calibrated thickness are perforated by xurography(17) and constrain the liquid height. Their assembly is simple and their geometry modular, thus allowing to create channels or wells that can fit the end-user application. (Fig.1B).

The micro-reactors are mounted onto the motorized stage of a microscope coupled with a DMD-based widefield illuminator that shines spatially modulated UV light. The DMD (digital micro-mirror device) allows arbitrary forms and gray levels (via pulse width modulation) to be projected. The radical generation rate is thus controlled in space and time (Fig.1C).

Overall the complete platform locally tunes the reagent concentration, the gas content and the UV-power. This control over the chemical context enables four hydrogel engineering operations with immediate applications in cell culture. We now describe these operations starting with total polymerization of hydrogel channels under inert atmosphere.

### Total polymerization

Perfusion channels and chemical gradients are critical to provide cells with essential nutrients and differentiation factors during growth. To this end we aimed at building walls and permeable barriers of arbitrary shape (movieS1) in our micro-reactors.

Yet, photopolymerization of hydrogels is impaired at the vicinity of PDMS as the perfusing oxygen forms an inhibition layer (deadzone)(18, 19). By generating an inert gas atmosphere over our microreactors, we deplete oxygen from the PDMS roof.

In such anoxic condition, the deadzone disappears and the cured structure reaches the ceiling forming a wall in the chamber (Fig.2A). Successive illumination patterns can be aligned in order to generate hydrogel-based millimeter-sized circuits (movieS2). Capillary driven flow is achieved by deposition of liquid drops at the micro-reactor’s entries. The perfusion of a solution of 1um fluorescent beads inside a hydrogel spiral showed no leakage demonstrating the total polymerization (Fig.2C, movieS2).

**Fig. 2.**
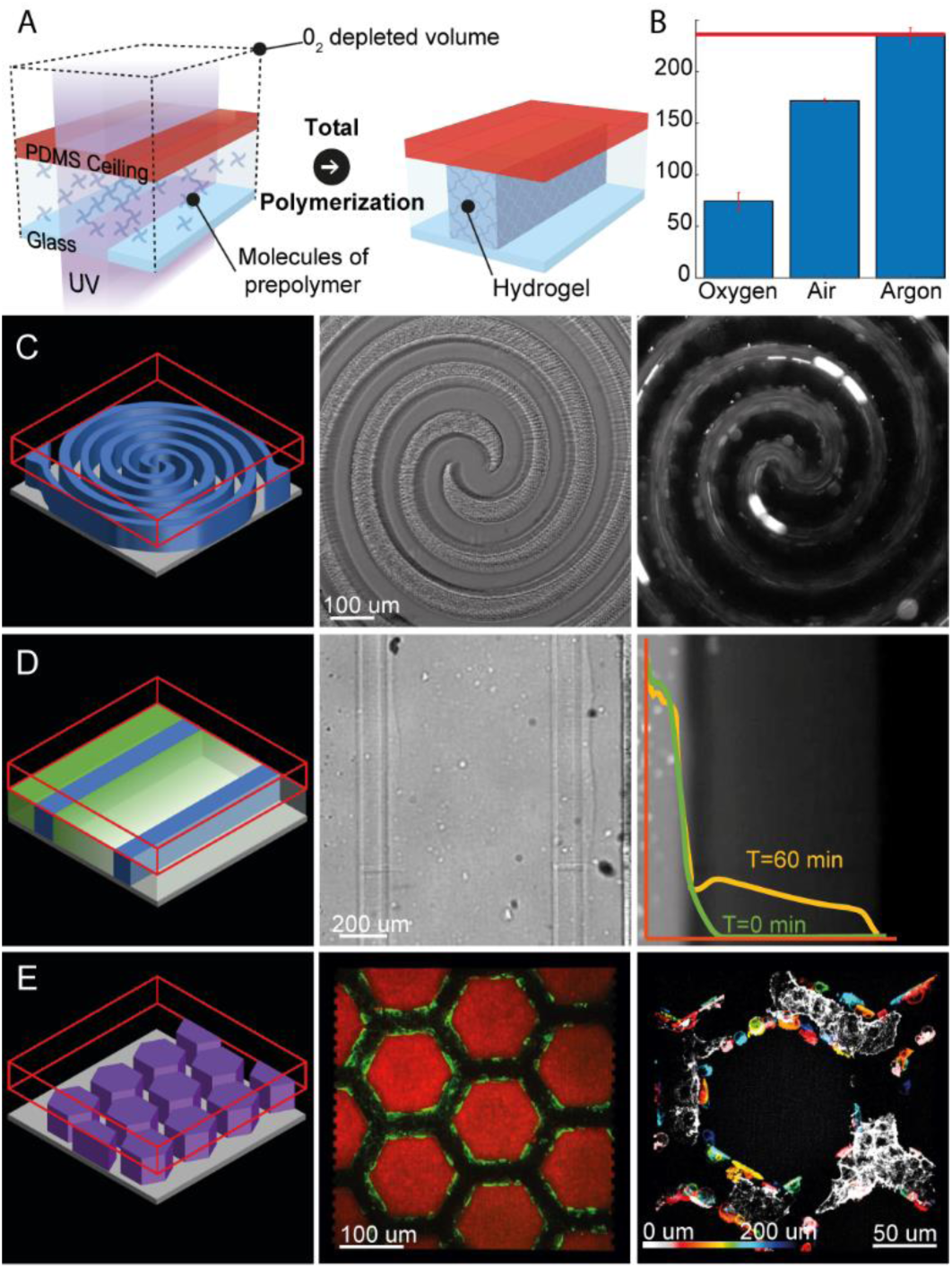
Building perfusable hydrogel channels with total-polymerization. (A), Schematics of the total polymerization procedure: hydrogel precursors (blue 4-branched stars) and photo-initiators are incubated. Argon perfusion generates an oxygen depleted volume (black dotted lines). After UV illumination (purple, purple arrows) the gel is polymerized(blue) in the whole insolated volume. Polymerized gel height (µm) is inversely correlated with the oxygen content, reaching the PDMS ceiling in its absence (B). Total polymerized hydrogels create leak tight barriers that support flow (C) and the generation of diffusive gradients (D). Biocompatible hydrogels can allow for the growth of cells in arbitrary vascular structures (E).

Secondly, hydrogel-channels can be tailored into semi-permeable membranes, letting small molecules pass through (Fig.2D). As an example, solution of purified GFP was let to perfuse through a photo-polymerized barrier, showing the establishment of a diffusing gradient over time (Fig.2D).

To turn these microfluidic channels into cellularized vasculatures, we complemented bioinert PEG hydrogels with cell adhesion molecules (Poly-L-Lysine, fibronectin). Hexagonal networks of such materials were entirely polymerized and used as cell culture templates. COS-7 cells colonized the hydrogel walls as revealed by 3D fluorescence imaging of the labelled cells after 3 days in culture (Fig.2E).

Additionally, we observed that the height of the hydrogel depends on the oxygen content of the atmosphere (Fig.2B). This can be used to globally control the thickness. However, the much-needed topographical structures, favoring spheroid organization or mimicking the complexity of organs, require local thickness control. Hereafter, we introduce a second engineering operation suited for the above purpose.

### Z-Controlled Polymerization

At a given oxygen content, the deadzone thickness is inversely correlated with the photon-flux(18). In our set-up, under air atmosphere, the PDMS ceiling simply imposes a constant oxygen boundary condition.

In turn, varying the laser power or the gray level intensity from 20 to 140 mW/cm^2^ dictates the resulting gel thickness from 65 to 175 µm respectively (Fig.3B).

**Fig. 3.**
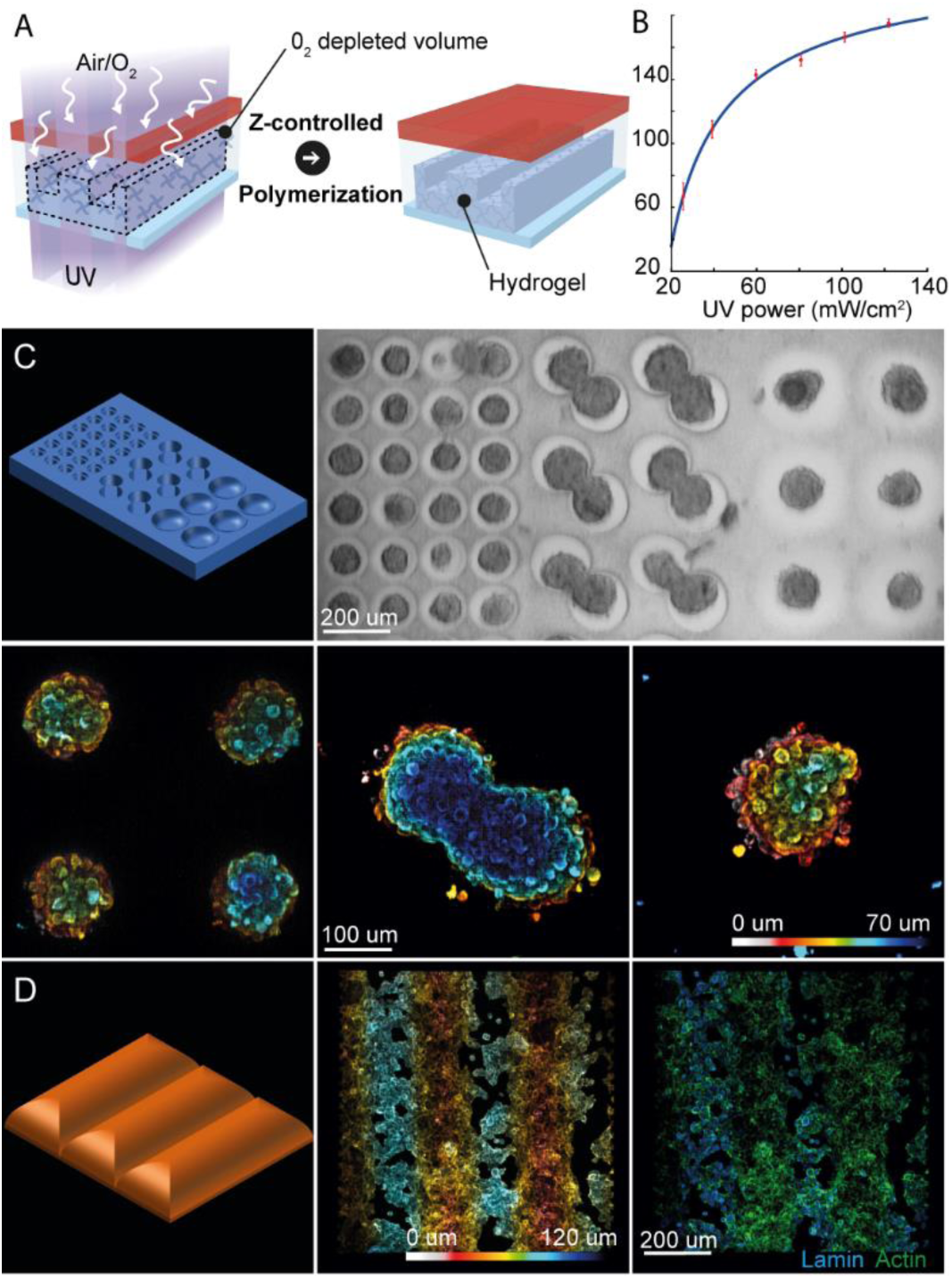
Controlling the gel topography with z-controlled polymerization. (A) Schematics of the z-controlled polymerization procedure: hydrogel precursors (blue 4-branched stars) and photo-initiators are incubated, air serves as an oxygen source. UV illumination (purple, purple arrows) generates an oxygen depleted volume (black dotted lines) with features corresponding to the photon flux distribution (darker and lighter purples). The polymerized gel (blue) will correspond to the oxygen depleted volume during the insolation. The gel thickness(µm) correlates with the local photon flux (mW/cm^2^) (B). Using this principle, grayscale pattern can generate topographical structures out of bioinert hydrogels to grow standardized spheroids (C). Alternatively the surface of biocompatible hydrogels can be colonized with cells (D).

Taking advantage of this structuration mode we created cup-shaped PEG templates ideal for spheroid culture (Fig.3C). Spheroïds are cell aggregates which can be readily used as physiological models or serve as a starting point for complex organoïds. Yet the lack of adequate culture substrates induces a high variability in shape and cell count leading to unreliable results.

Here, HEK cells seeded on topographically structured PEGs aggregated into standardized spheroids the size and shape of which coincided with the features of the micro-wells (Fig.3C).

The inertness, transparency, permeability and simple structuration of this hydrogel make it the go-to material to dictate spheroid growth.

When needed, the inertness of the PEG can be reverted to allow for cells to colonize the topographies. Here, cell-adherent waves were created by mixing Matrigel with a photopolymerizable PEG precursor. Matrigel is a temperature-curing hydrogel enriched in laminin, collagen and other adhesion factors ensuring high cellular compatibility. On these hybrid hydrogels, seeded cells quickly spread homogeneously, forming a confluent layer on top of the structure (Fig.3D).

Even in the absence of photopolymerizable precursors, some prepolymers can undergo total and z-controlled polymerization, as shown for Matrigel (MovieS3). However, UV-induced crosslinking drastically alters the gel rheological properties which are critical for cell behavior(20). To overcome this limitation, we introduce a third engineering operation where most hydrogels can be structured without significantly altering their final mechanical properties.

### Photo-scission

Photo-scission removes areas of hydrogels while the remaining mesh is left unscathed. While oxygen is known to inhibit radical photo-polymerization and induce the photo-scission of PEG chains, the latter reaction has found only few applications in material science(21). In this report, we show how photo-scission unlocks the structuration of bulk, native hydrogel such as PEG, Poly-acrylamide, Agar-agar and Matrigel. (MovieS4).

This reaction occurs when a cured gel, perfused with photo-initiators, is exposed to UV light in the presence of a sustained supply of oxygen. Indeed, anoxic conditions inside the insolated volume would favor crosslinking instead of chain scission. Here again, a permeable PDMS ceiling or floor (Fig.4A) is in contact with the air atmosphere thus providing a continuous oxygen supply during the reaction. The gel will then progressively liquefy in material-specific kinetics (Fig.4B).

**Fig. 4.**
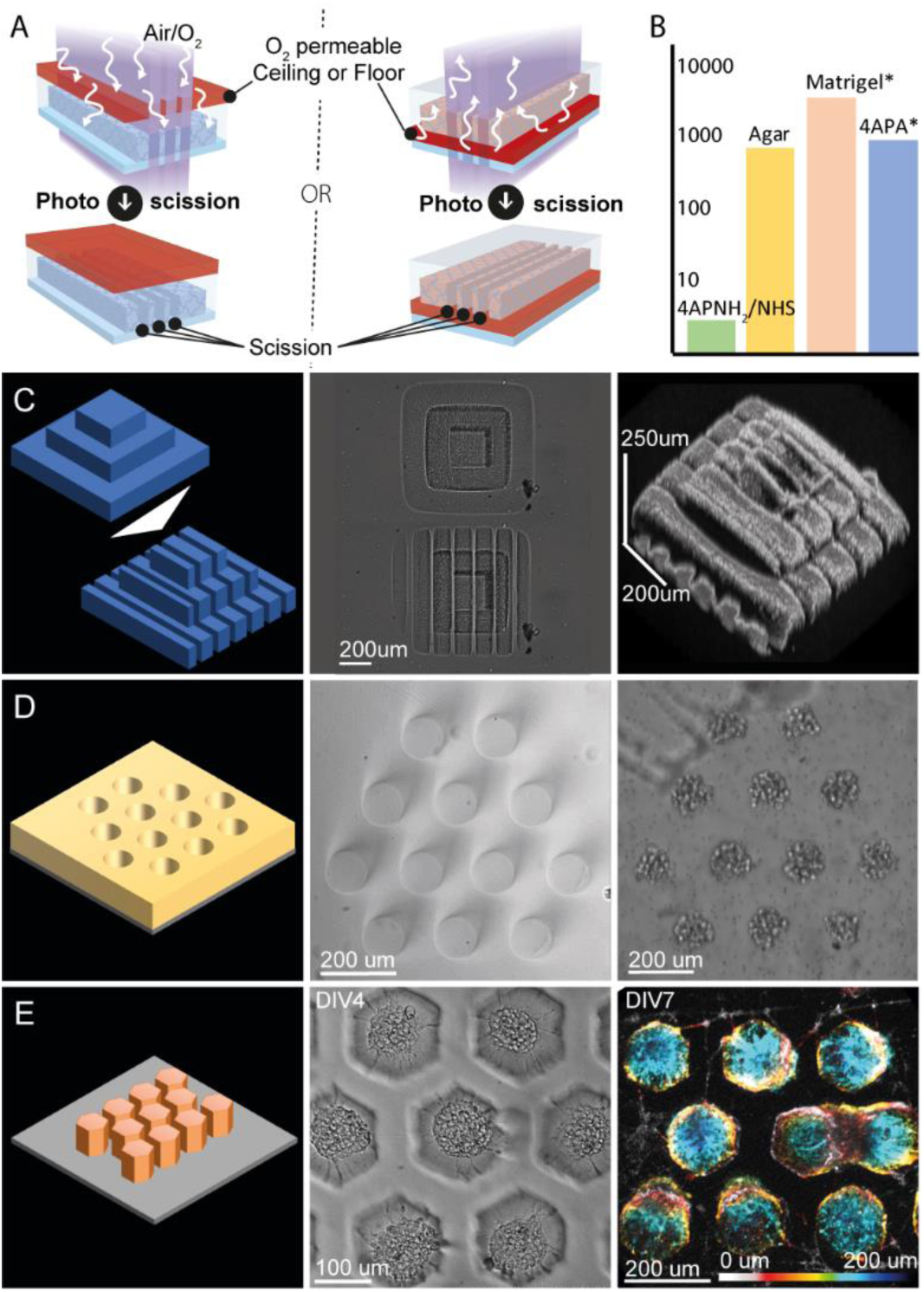
Subtractive manufacturing of common hydrogels with photo-scission. (A) Schematics of the photo-scission procedure: cured hydrogels (blue, and pink) are exposed to patterned UV irradiation (purple, purple arrows) in the presence of photo-initiators. The PDMS ceiling or floor (red) ensures that oxygen is renewed in the insolated areas. This ultimately leads to the local liquefaction of the hydrogel. For the same pattern, scission time (s) is material dependent and may require dilution of the photo-initiator (*) (B, logarithmic scale). Photopolymerized hydrogels can also undergo photo scission (C). Soft bio-inert agar can be photo-scissionned to form spheroid assembly templates (D). Biocompatible pillars of Matrigel can be created to grow neurons (E).

Soft, visco-elastic hydrogels such has Matrigel are widespread but lack simple structuration means. Indeed, the replica molding of such frail material is notoriously difficult. Here, we exemplify our contactless photo-scission on very soft materials (Matrigel, Agar 0,5%), to engineer high aspect ratio (>2) structures.

An agar-agar template consisting of an array of 100um diameter microwells served as a spheroid culture substrate. (Fig.4D)

With the same method, hexagonal pillars were carved out of a Matrigel layer. These structures were among the softest we created (400 Pa) and were especially suited for neurons cell culture. As the pictures demonstrates, the neurons soma aggregated on top of the pillars while the axons spread out (Fig.4E) forming interconnected spontaneously firing networks after a week (MovieS5). Of note photo-polymerized hydrogels can be subsequently photo-scissioned. In this example, a photo-polymerized “Aztec temple” was generated using the two previously described engineering operations: a square pillar was total-polymerized while two stages were added by z-controlled polymerization. The structure was then sliced demonstrating that cross-linked objects can be photo-scissionned (Fig.4C, MovieS6)

So far, the structures we’ve described were either cell repellant or fully adhesive. In the latter part we’ll show how to decorate (locally functionalize) inert hydrogels with adhesive biomolecules.

### Hydrogel Decoration

PEG or agarose hydrogels are bio-inert: they prevent biomolecules from adsorbing and thus cells from adhering(22). Previously, they have been either homogeneously(3) or locally(9) complemented with biomolecules to promote cell adhesion and differentiation. Photo-linkers like Sulfo-Sanpah or Acryl-PEG-Sva (APSv) are a family of commercially-available molecules capable of hydrogel decoration. One extremity of these hetero-bifunctional molecules can photo-graft to a hydrogel while the other moiety can bind amine-rich adhesion molecules like poly-L-lysin via esterification.

Here, we attach an acryl-PEG-Sva linker onto a PEG hydrogel in a spatially controlled manner by the conjugated action of the acryl moiety, the photo-initiator, and the patterned UV illumination (Fig.5A).

**Fig. 5.**
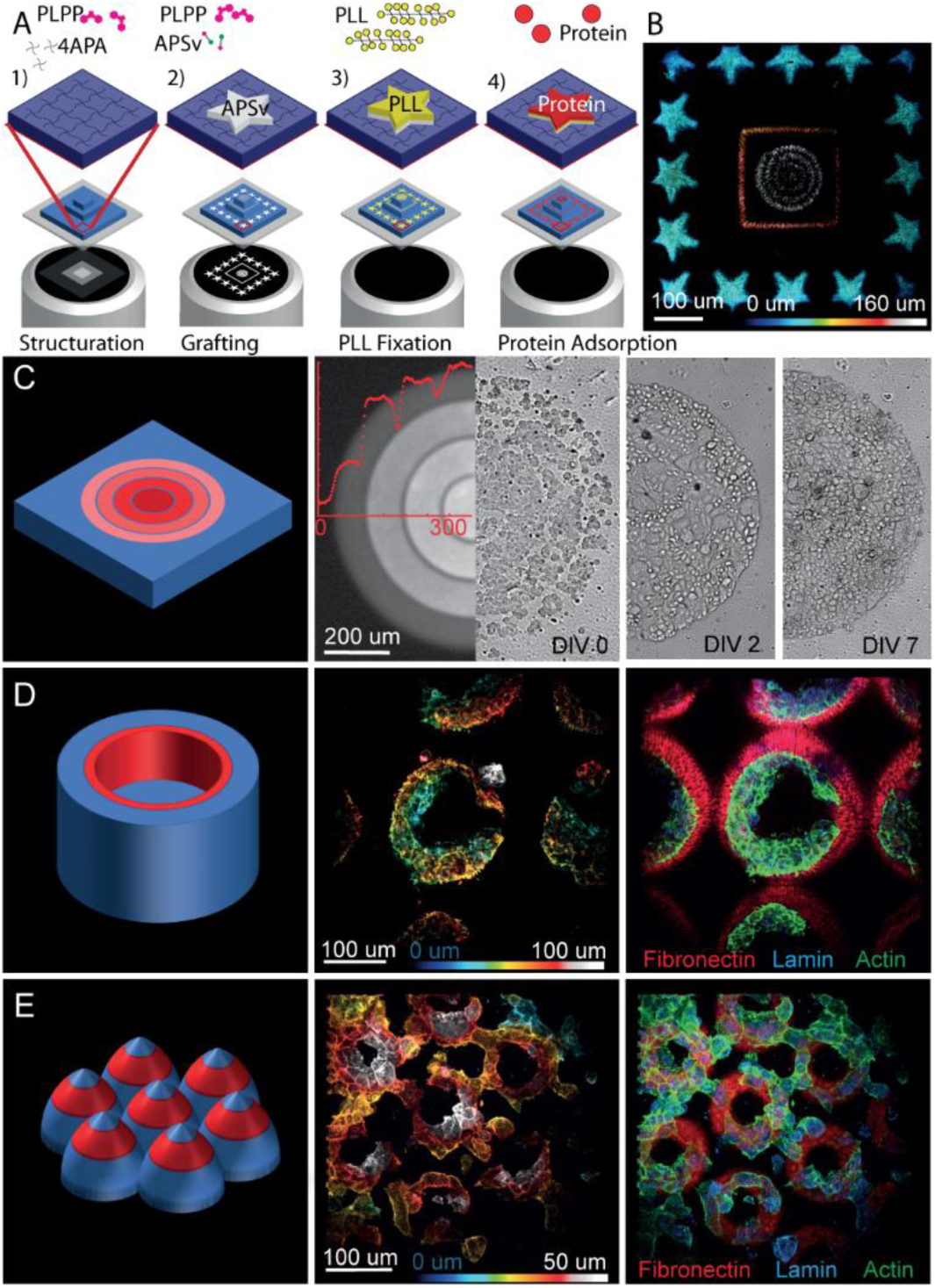
Decoration: quantitative and patterned functionalization of hydrogels. (A) Schematics of the micro-structuration and decoration of an “Aztec temple” hydrogel. First a gel is formed (dark blue) via height-controlled polymerization of 4-arm-PEG-acrylate (blue) with PLPP (pink). Then acryl-peg-sva (pink and green, white layer) is photo-grafted in the presence of PLPP according to the projected UV pattern. This allows covalent ester-binding of poly-L-lysin (yellow) and the subsequent adsorption of biomolecules (red). Complex decoration patterns can be easily aligned on previously generated topographies (B). Additionally, the process is dose dependent and can create adhesive gradients (C). It is thus possible to cultivate cells in microwells (D) or topographies (E) that combine adhesive and structural cues.

The photo-grafted succinimide-ester will subsequently react with the many amino-groups of the poly-L-lysin. This locally reverses the gel antifouling properties and permits subsequent adsorption of a wide range of proteins (23).

The photo-addition of acryl-PEG-Sva is UV dose-dependent and as such the density of grafted poly-L-lysin can be modulated. In Fig.5C, 4 concentric levels of photo-linkers were grafted on a flat PEG hydrogel leading to differential covalent binding of poly-L-lysin-FITC. We were able to induce differential cell adhesion by adsorbing fibronectin on this pattern. Using the same approach, laminin, a neuron specific adhesion molecule, was successfully grafted to orient neuron growth (S1).

With this maskless decoration method we can easily align biomolecule motifs onto previously built topographies. As a proof of concept, a 3-stage Aztec temple was polymerized as described in Fig.4 and each stage was decorated with a distinct motif (Fig.5B).

We now demonstrate that our platform precisely controls cell positioning on tridimensional objects. First, PEG Hydrogels were polymerized with power and grayscale modulation to obtain topographies. They were subsequently decorated with the aforementioned method allowing COS-7 cells to colonize these structures reminiscent of capillaries (Fig.5D) and intestinal villi (Fig.5E).

In summary, we combined UV-patterned illumination with gas permeable reactors to obtain 4 hydrogel engineering operations: total polymerization, z-controlled polymerization, photo-scission and decoration. We highlighted that they can readily address hydrogel engineering challenges alone or in combination.

## Discussion

As any fast-growing field, 3D cell culture has known a burst in material and application diversity. Our platforms aim at embracing many materials (generic) and protocols (flexible) while being user-friendly (accessible). We’ll now discuss these three aspects with emphasis on structuration, decoration and the role of the DMD.

Together, our results suggest that the chemical reactions leading to hydrogel structuration are driven by oxygen, light and benzophenone regardless of the material. This is the root of our multi-material capability.

As a result, total and z-controlled polymerization do not require prepolymers with photo-polymerizable moieties (movieS3). Even though, acrylated polymers display a faster crosslinking rate, our benzophenone-based initiator is able to induce polymer photo-crosslinking by itself. We also observed that chain length is critical to achieve sharp structures with good shape replication fidelity (S2C).

Indeed, polymerization of short prepolymer is impossible for native molecules and uncontrollable with acryl-moieties probably due to their high diffusivity. In our hands, long branched acrylated prepolymers are ideal to quickly form sharp structures (S2C).

In contrast, photo-scission liquefies all the hydrogels we have encountered so far regardless of their chain length. Nevertheless, this oxygen-consuming process is slower than photo-polymerization. Oxygen starvation prevents photo-scission and leads to crosslinking at depleted points in the insolated volume. Higher laser power and photo-initiator concentration increase the scission rate. Nevertheless, for a given geometry they must be low enough to prevent oxygen depletion. Finally, the above parameters together with material specific kinetics lead to highly variable scission times. For instance, the scission of dense agar structures can take hours while it only takes seconds to obtain sparse PEG gel geometries.

In line with the multi-material capabilities of our structuration method, a decoration operation supporting a wide range of hydrogels and biomolecules was needed.

Fortunately, there are plenty of available linkers which can be chosen depending on the material. Photo-linkers such as Sulfo-SANPAH can be grafted onto most substrates while avoiding further gel crosslinking(24). However, they are moisture and light sensitive making them very unstable and thus dispendious. To cope with those limitations, we combine our light stable photo-initiator with accessible Acryl-PEG-Sva linkers. This provides a user-friendly alternative at the expense of unwanted crosslinking inside the insolated area.

The poly-L-lysin incubation step was implemented to increase to robustness and flexibility of the decoration. Indeed, the photo-linker’s succinimide ester hydrolyze in water limiting incubation time. Thus, optimal grafting would require highly concentrated biomolecules with many available amino groups but protein solutions are often highly diluted and can lack free amines. In contrast, poly-L-lysin quickly grafts onto the succinimide due to its high number of free amines. This now-stable grafting intermediate allows for the enrichment of most biomolecules even when highly diluted or poorly reactive.

We previously emphasized that our platform was suitable for structuring and decorating many hydrogels each having its own reaction rate. Here we introduce patterned light as a straightforward method to locally control the kinetics of the photoreactions that occur in the micro-reactor.

The DMD is the only hardware capable of locally tuning the UV photon flux within an entire illumination field. In essence, a DMD is a binary light modulator, the mirrors can only flip between ON and OFF state. It interprets grayscales by quickly switching between those two states in a process called pulse width modulation (PWM). The flipping rate of the mirrors is fortunately faster than the reaction kinetics so that PWM can control the thickness as efficiently as the laser intensity (S2A). This gives us local control over the photon flux. We also implemented an alternative insolation mode which interprets gray levels as UV doses. In this scenario, each pixel has a continuous insolation time proportional to the gray level. Together with the tunable laser power, this gives us enough flexibility to quickly optimize the illumination settings leading to the desired template.

We previously demonstrated that UV dose was instrumental in quantitative decoration and the photon flux was key to the generation of topographical structures. To give an insight into the possibilities offered by these illumination modes, one can fix the overall thickness with the laser power and locally tune the crosslink density with the UV dose(10). Hydrogel structures with orthogonal structural and rheological features are then feasible (S2D).

Lateral diffusion of oxygen during polymerization makes patterns with fine features especially difficult to print. The resulting structures appear as trimmed on the sides. To overcome this limitation, we optimized patterns design by offsetting the “zero” to a low value (S1B). This generates a photon flux sufficient to induce oxygen consumption without initiating polymerization ultimately reducing the trimming effect.

To summarize, the illumination set-up turned multiple 4-dimensional reaction kinetics into a playground where simple grayscale images sets the rules of the desired hydrogel micro-niches.

### Outlook

Engineering soft gels for 3D cell culture is usually a difficult task. Here, by combining gas-permeable reactors together with patterned UV illumination, we ended up with a flexible yet simple hydrogel design solution:

When applied to its fullest, this principle can make perfused systems in short time scales. It can also generate topographically complex structures in one step. It gives new structuration means for photo-inert hydrogels. It can decorate gels to promote cell adhesion. Each of the engineering operations above can be queued since alignment steps are easy.

As biologists are striving to create standardized and physiologically relevant in vitro models using many materials, cells and protocols, our set-up can address many of their challenges. Overall, this platform has the potential to offer a common frame work for the scientists to share and expand their 3D cell culture discoveries.

## Acknowledgments

The authors thank Nikon Instruments France for loaning some of their equipment. We thank our readers on bioRxiv for their comments and ideas in order to improve the impact of our work. We thank Corey Butler for careful reading of the manuscript.

## Author Contributions

AP conducted the experiments. All authors designed the experiments and wrote the manuscript.

## Methods

### Experimental Setup

For all the experiments, we used a Nikon TI-E inverted microscope (Nikon Instruments) equipped with a DMD-based UV patterned illumination device (PRIMO, Alveole, France) together with its dedicated software (LEONARDO V3.3, Alveole). The wavelength of the illumination laser is 375 nm. 4X S-Fluor and 10X Plan Fluor (Nikon) were used to ensure efficient UV transmission. A homemade version of this illumination device was previously described in 21. The patterns projected with PRIMO for all the experiments can be found in S4.

### Modular Microreactors

We used custom PDMS chips made of two superposed PDMS thin films with holes fabricated by xurography (PDMS Stencils, Alveole, France, previously described in 21). These stencils were stacked onto a 22×22 mm 170 um thick glass coverslip (Schott Nexterion, Schott Jena, Germany) forming PDMS microchambers of various geometries (Fig.1). Depending on the application, the upper PDMS film was removed after the building steps to ease the seeding of cells or beads. In gas-controlled structuration experiments, the device is placed in a custom enclosure allowing argon or oxygen perfusion during the insolation steps.

### Reagents were purchased as follows

4-Arm-PEG-Acrylate MW 10K, Acryl-PEG-SVA MW 2K, 4-Arm-PEG-Amine MW 10K, 4-Arm-PEG-SAS MW 10K. Laysan Bio (Arab, USA). Matrigel Corning (USA). PLPP 1X (14.6 mg/mL) (Alvéole, France). Human Fibronectin (Roche). Agar low melting point, PEG 8K, PEG 20K, PEGDA MW700, Laminin from mouse EHS, PolyLysine, PolyLysine FITC conjugate (P3543), NH4Cl, Triton X100, Paraformaldehyde, Bovine Serum Albumin (BSA), Phosphate Buffered Saline (PBS), N,N-DimethylFormamide (DMF), Dulbecco Modified Eagle Medium, NeuroBasal (Sigma Aldrich). 0.3 um diameter carboxy-fluorescent latex beads, Phalloidin-Alexa647 conjugate (Invitrogen), anti-Lamin (Abcam, ab16048) and Goat Anti-rabbit antibody Alexa 568 conjugate (Abcam, ab175471). Sir-Actin (Spirochrome), Fluo4AM (Thermo Fisher), Membrite (Biotium).

### Total Polymerization

precursor gel solutions containing monomer and PLPP as photo-initiator were incubated inside the PDMS microreactors. Under inert atmosphere (argon, N2), gels were polymerized using maximal laser intensity. The insolation times varied from 20sec to 5 minutes depending on the monomer polymerization yield. Precursor gel solutions used in the article were as follow: 50 mg/mL 4-arm-PEG-Acrylate diluted in 1X PLPP (Fig.2B, C, D, MovieS2, MovieS6), 25 mg/mL 4-Arm-PEG-acrylate with 0.5 mg/mL of Poly-L-Lysine in 0.5X PLPP (Fig.2E, MovieS1). To functionalize the hydrogels in Fig 2E, 100 ug/mL of Fibronectin were incubated for 15 minutes prior to cell seeding.

### Z-controlled / Topographical Polymerization

Precursor gel solutions containing monomer and PLPP as photo-initiator were incubated inside the PDMS microreactors. Gels were polymerized in the presence of air using a combination of grayscale projection and/or variable light intensity. In one mode a gray scale pattern is insolated with gray levels interpreted as photon fluxes leading to a topographical structure. In another mode the gray levels are interpreted as UV dose and the laser intensity is set to obtain the desired thickness. In this second mode multiple projections must be done to obtain topographies. The insolation times varies from 20sec for 4 arm-PEG-acrylate to 5 minutes for Matrigel alone depending on the monomer polymerization yield and photon flux. Precursor gel solutions used in the article were as follow: 50 mg/mL 4-arm-PEG-Acrylate diluted in 1X PLPP (Fig.3B, C, S2A, B, D, E, MovieS6), Matrigel 5mg/mL in 0.5X PLPP (MovieS5),1.250 mg/mL 4-Arm-PEG-Acrylate with 5mg/mL Matrigel in 0.75X PLPP (Fig.3D). 100 mg/mL PEG 8K, 100 mg/mL PEG 20K, 100 mg/mL 4-arm-PEG-Acrylate and 100 mg/mL PEGDA 700 all in 0.5X PLPP (S2C).

### Photo-scission

Hydrogel precursor solutions were allowed to cure inside the PDMS microreactors. PLPP was left to perfuse in the hydrogels for 10 minutes prior to UV projection. The PLPP concentration was also adjusted depending on the material: 0.0125X for Matrigel or 4-arm-PEG-acryl and 1X for agar-agar or 4-arm-PEG-amine / 4-arm-PEG-SAS. Photo-scissionned hydrogels in the article were the following: 50 mg/mL 4-Arm-PEG-Acrylate (Fig.4C, MovieS3, MovieS6), 20 mg/mL 4-Arm-PEG-Acrylate(Fig.4B), 20 mg/mL Agar (Fig.4B),0.50 mg/mL Agar (Fig.4D, MovieS3) 20 mg/mL Poly-Acryl-Amide (MovieS3), 20 mg/mL 4-Arm-PEG-Amine /4-Arm-Peg SAS (Fig.4B, MovieS3), 21mg/mL Matrigel (Fig.4B,E, MovieS3)

### Decoration

Solutions consisting of 50 mg/mL 4-arm-PEG-acrylate in 1X PLPP were photopolymerized with z-control inside the PDMS microreactors. For agar and 4-arm-PEG-amine/4-arm-PEG-SAS (S3) solutions of 20mg/mL of precursor were let to cure in the microreactors. After rinsing with PBS, a solution of 100mg/mL of APSV in 1X PLPP was incubated. These solutions were prepared immediately before use. 1g/mL aliquots of APSV in DMF can alternatively be used, they are stable for 2 months at -20C°. The photo-linkers were grafted onto the surface of the gel by the subsequent insolation which last for 30 sec to 2 minutes depending on photon flux and substrate. The gels were rinsed and incubated with 1mg/mL PLL immediately after the insolation step to prevent the activated esters to hydrolyze. After one hour of PLL incubation the gels were rinsed. To promote cell adhesion, 100 ug/mL of Fibronectin (Fig.4) or 25ug/mL laminin (S1) were incubated for 15 minutes prior to cell seeding.

### Cell Culture

COS-7 and HEK 293T cells were seeded and cultivated in complete cell culture media (DMEM, FBS). E18 rat primary cortical neurons were seeded and cultivated in complemented neurobasal.

### Staining and imaging

Fixation and staining of cells was conducted using standard procedures. Staining of gels structures was performed by coating with 0.3 um carboxy-fluorescent beads. Neurons on Matrigel (Fig.4E MovieS5) were imaged using Sir-Actin and Fluo4AM live markers. Spheroids (Fig.3C) were stained using membrite live marker. 3D imaging was achieved using a homemade DMD based confocal microscope(25).

## SI APPENDIX

### Supplementary Figures

**S1.**
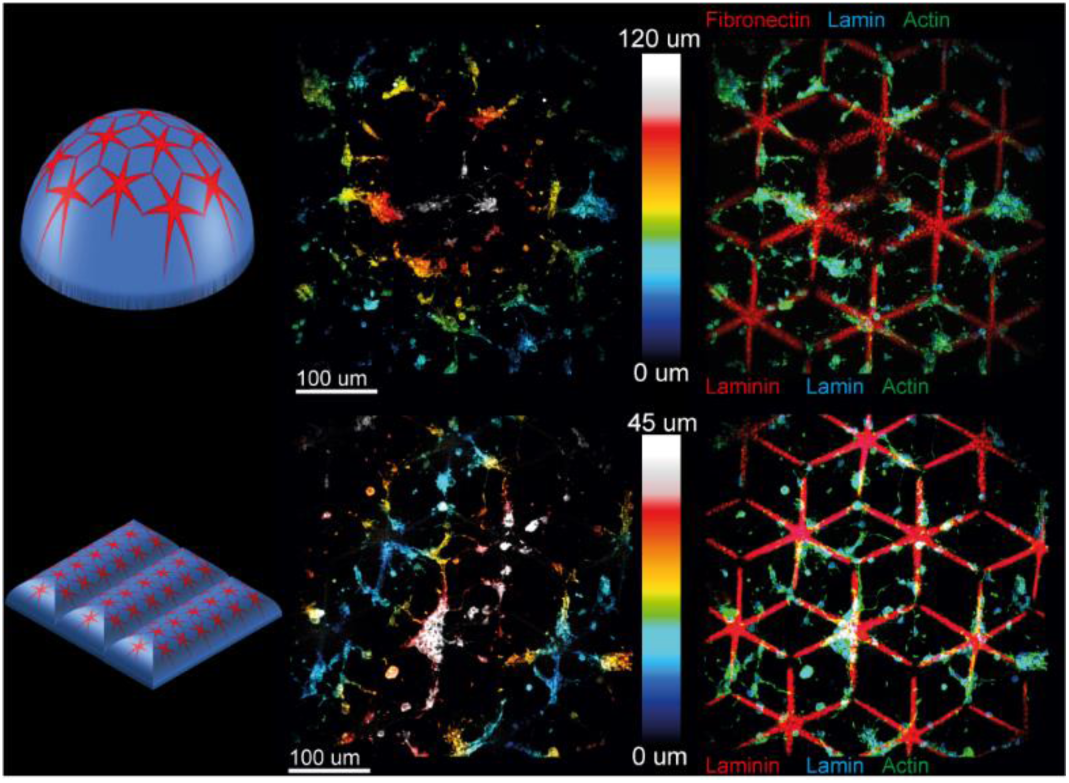
Decoration of hydrogels for neurons. left panels: schematic representation of 4-arm-PEG-hydrogels (blue) decorated with Poly-L-lysin(red) and laminin. Middle panels: Z-color-coded confocal fluorescence microscopy images of neurons seeded on the gels and stained for actin. Right panels: 3-color max-projection of confocal microscopy z-stack showing the patterned molecules (red), the actin cytoskeleton (green) and the nuclear envelope (blue).

**S2.**
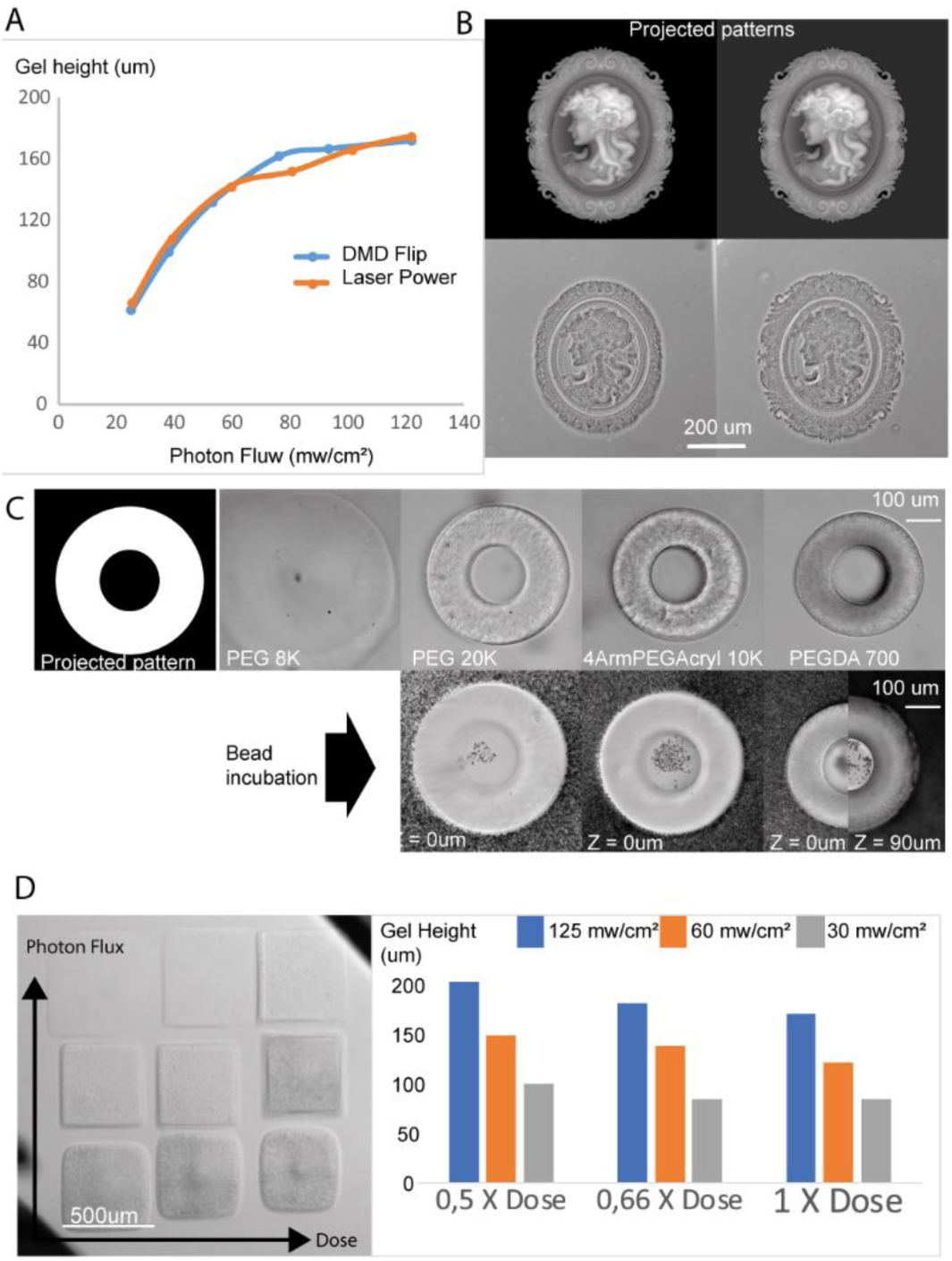
Optimization of photo-polymerization. The gel height is equally modulated by the laser power or the projected gray levels (A). A gray background can nullify oxygen trimming (B). Polymerization fidelity depends on precursor length (C). Gel thickness and reticulation are decorrelated (D).

**S3.**
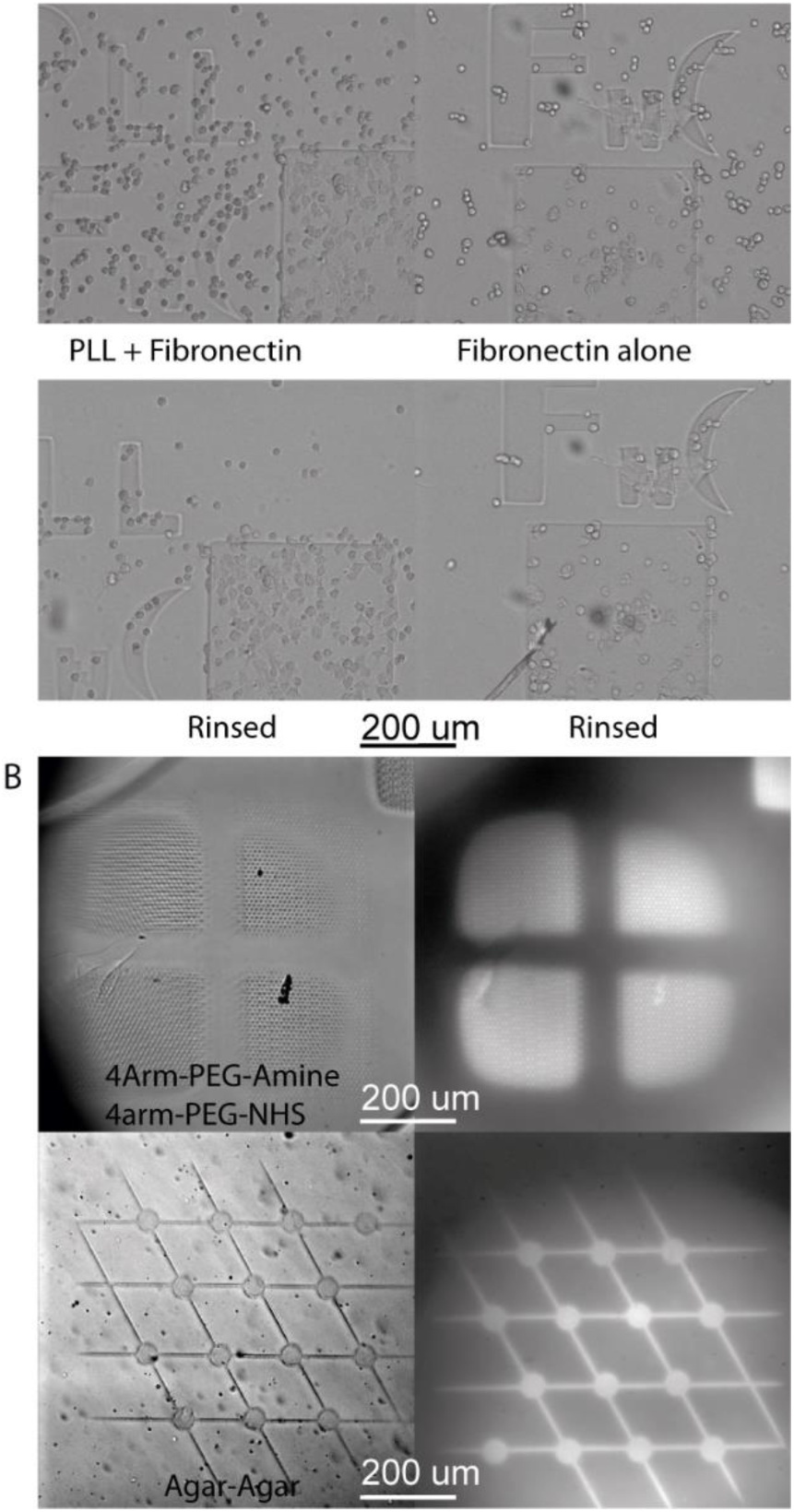
Optimization of decoration. The addition of poly-L-lysin increases cell adhesion (A). Non acryls hydrogels can be decorated (B).

**S4.**
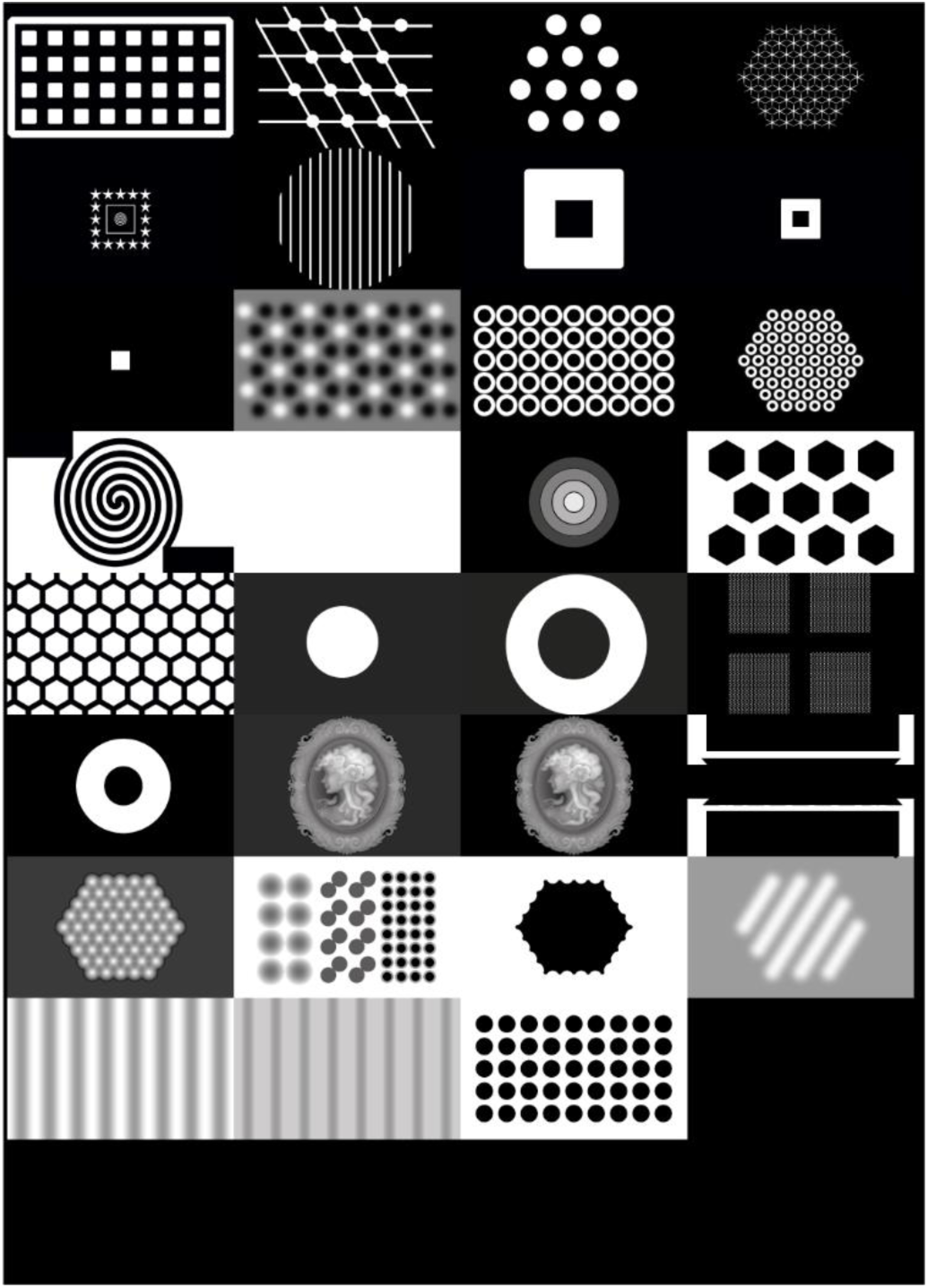
All the patterns used in this report.

### Supplementary Movies

MovieS1: Time-lapse showing how total polymerized 4-arm-PEG-acrylate mixed with poly-L-lysin swell and shrink due to osmotic pressure under successive rinsing with de-ionized water and PBS.

MovieS2: Total polymerization steps can be queued to create hydrogel-based fluidic channels that support flow. left panel: schematic view showing micro-reactor design and polymerized hydrogels. right panel: time-lapse of the fabrication.

MovieS3: First part: time-lapse showing how the “PLPP” photo-initiator can initiate the photopolymerization of non-acryl prepolymers like Matrigel. Second part: Z stack demonstrating the control of the thickness by tuning the photon flux as shown by the deposition of beads.

MovieS4: Time-lapses highlighting the photo-scission of various hydrogels materials: from left to right 4-Arm-PEG-acrylate, 4-Arm-PEG-NH2/NHS, Poly-Acryl-Amide, Agar-Agar, Matrigel.

MovieS5: Calcium imaging of neurons growing on top of Matrigel pillars (7 days in vitro) highlighting spontaneous neuronal discharges (fluorescence spikes).

MovieS6: Time-lapse demonstrating the successive structuration step queued. First part: total polymerization of a pillar. Second part: z-controlled polymerizations of two additional stages. Third part: photo-scission of small lines to cut the “temple” in slices.

